# Electroencephalographic responses before, during, and after upper limb paired associative stimulation

**DOI:** 10.1101/2025.01.29.635408

**Authors:** Yumi Shikauchi, Kazumasa Uehara, Yuka O. Okazaki, Keiichi Kitajo

## Abstract

Paired associative stimulation (PAS) is a non-invasive protocol involving repeated stimulus pairs to activate two cortical areas alternately, inducing Hebbian-like plasticity. However, its neurophysiological impacts remain unclear. To determine the changes that occur in the brain during PAS, brain activity during PAS must be measured and distinguished from the electromagnetic artifacts produced by the stimulation. Here, we present a novel dataset of electroencephalography (EEG) measurements during PAS with an inter-stimulus-interval of 25 ms (PAS_25_, expected to induce long-term potentiation-like changes) or 35 ms (PAS_35_, no expected change). This dataset includes raw data and pre-processed data with electromagnetic artefacts removed. he right ulnar nerve’s electrical stimulation preceded transcranial magnetic stimulation to the left primary motor cortex in both cases. EEG was measured before and after the PAS sessions, with only electrical or magnetic stimulation. To demonstrate the quality of the data, we summarize the stability of the stimulation site and the event-related potentials before, during, and after PAS. This dataset will enable observing brain dynamics due to the accumulation of stimulations during PAS and differences in responsiveness to stimulations before and after PAS.

## Background & Summary

Paired associative stimulation (PAS) is a unique paradigm for inducing Hebbian-like plasticity in the intact human brain^1,2^. It was first demonstrated that transcranial magnetic stimulation (TMS) of the primary motor cortex (M1) after peripheral electrical stimulation of the median nerves increases the amplitudes of motor-evoked potentials (MEPs) in the abductor pollicis brevis muscle^2^. Subsequent research has shown that repeated associated stimulation of two cortical sites at an appropriate inter-stimulus interval (ISI) can alter not only MEP amplitude but also functional connectivity and cognitive motor performance^3–7^. PAS is a powerful paradigm that can be applied to study mechanisms of cognitive functions and their underlying functional connectivity, as well as diseases involving neural plasticity. However, its efficacy varies widely among individuals^8^. When electrical stimulation is delivered to the upper limb as a preceding stimulus 21–25 ms before TMS to M1^4^, 39–75% of participants show a long-term potentiation (LTP)-like response^8–10^. The relative timing between synaptic input and postsynaptic activity modulates synaptic strength, as described by Hebb’s rule. Delivering TMS to M1 25 ms after electrical stimulation of the arm represents an optimal relative timing to induce LTP-like changes in S1-M1 coupling. In contrast, a longer interval between stimuli (e.g., 35 ms) no longer constitutes a relative timing that influences the plasticity of synaptic strength, resulting in no change in coupling^1,11^. Generally, the responsiveness to TMS, such as the amplitude of TMS-evoked potentials (TEPs) and MEPs, depends on dynamic brain states (e.g., the phase of brain oscillatory activities)^12–15^. Conversely, it has been suggested that cortical thickness and M1 excitability explain the differences in the effects of PAS between responders and non-responders^8,9,16,17^. However, state-dependency in brain dynamics has not been demonstrated. Simultaneous measurement of PAS and electroencephalography (PAS-EEG) is essential to examine functional changes in the brain during PAS. Although the simultaneous measurement of TMS and EEG (TMS-EEG) has become possible due to recent advances in amplifiers, there are still many technical challenges^18^. Only a few research teams have been successful, and thus sharing TMS-EEG data is extremely important^19–21^. To our knowledge, no public datasets are available to study brain activity changes during PAS (i.e., PAS-EEG).

There are two major technical issues in measuring brain activity by scalp EEG during PAS. First, the placement of the TMS coil is obstructed by the measurement device (e.g., EEG electrodes and lead wires), which results in a longer distance between the TMS coil and targeted brain region. This leads to less effective and localized TMS effects^22^. Second, the monophasic TMS pulses employed in the PAS protocol result in higher motor thresholds and limited cortical effects compared with biphasic pulses used in other paradigms^23^, thereby requiring higher TMS intensities to induce PAS effects. Along with the stimulus intensity, the EEG signal is severely contaminated by various stimulation-derived electromagnetic and physiological artifacts, making it difficult to obtain high-quality EEG signals in the PAS-EEG paradigm^24–27^.

Here, we present a dataset of PAS-EEG experiments conducted under two conditions in two separate day sessions: (1) ISI = 25 ms (PAS_25_), where a long-term potentiation-like change is expected and (2) ISI = 35 ms (PAS_35_), where no change is expected^1^. These conditions enable verification of the time-dependent effects of PAS. In both conditions, electrical stimulation of the right ulnar nerve preceded TMS to the left M1. Before and after the PAS sessions, EEG measurements were also taken with electrical stimulation alone (ES sessions) and magnetic stimulation alone (TMS sessions) (Fig. 1). All experiments were conducted in a double-blind fashion, with both the experimenter and participants unaware of the condition to which they were assigned. During stimulation, EEG signals were recorded from nine healthy volunteers using the 10-10 international system, which involved 63 scalp sites via flat Ag/AgCl TMS-compatible electrodes. To ensure the quality of our data, we summarize the event-related potentials before, during, and after PAS. In recent years, methods that actively mask the auditory co-stimulation associated with TMS by presenting appropriate noise simultaneously with TMS have become popular^28–31^. However, in this study, we only used earplugs to passively block the clicking sound, as there are no reports on the effects of auditory stimulation during PAS. This dataset will make it possible to observe brain dynamics due to the accumulation of stimulations during PAS and differences in responsiveness to stimulations before and after PAS.

**Figure 1.**
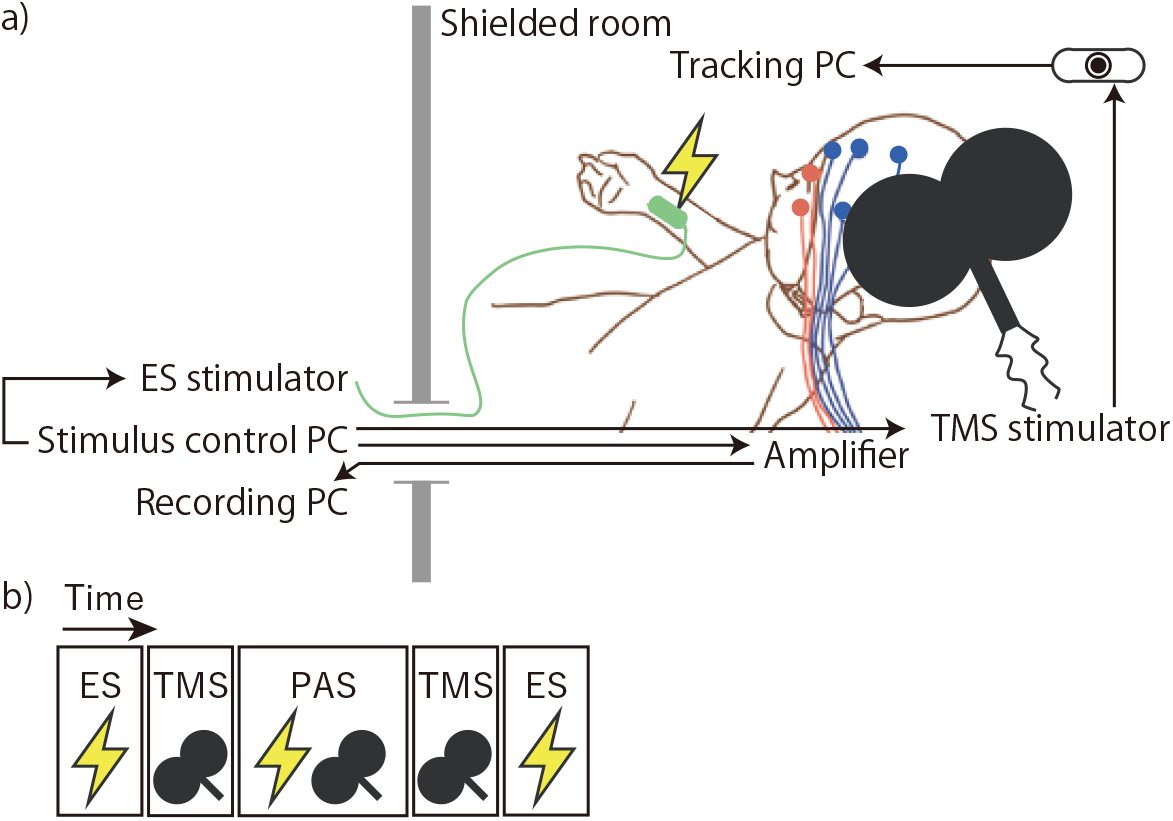
Schematic representation of the experimental procedure. a) The set-up. Blue and pink circles indicate electroencephalography and electrooculography channels, respectively (for clarity, not all of them have been depicted). The oval object in the upper right corner is a tracking camera that monitors the TMS posture. b) Sessions with only electrical stimulation (ES) and only transcranial magnetic stimulation (TMS) were conducted before and after paired associative stimulation (PAS).

## Methods

### Participants

We recruited nine healthy volunteers (four women and five men, aged 20–36 years, mean age: 26.8 ± 5.2 [SD] years) who provided written informed consent before participating in this study. The inclusion criteria were as follows: (i) aged 20–39 years; (ii) right-handed; (iii) no history of neurological disorders including seizure, febrile convulsion, or phobia; and (iv) not currently taking any psychotropic medication. Participants were screened for contraindications to TMS according to established safety guidelines^32,33^ and were consistent right-handers according to the Edinburgh Handedness Inventory^34^. The RIKEN ethics committee approved this study, which was conducted following the Declaration of Helsinki and according to institutional regulations based on research guidelines for the COVID-19 pandemic^35^.

Due to interruptions caused by the COVID-19 pandemic, we were unable to achieve the initially planned sample size. Additionally, data from two participants were excluded from analysis due to insufficient data quality. As a result, we obtained usable data from seven participants for analysis.

### Experimental paradigm

Each participant attended two double-blind PAS sessions separated by a week. We conducted all sessions in the afternoon to avoid any effects of circadian modulation of cortical suppression^36,37^. In these PAS sessions, the ISIs for peripheral nerve ES and TMS were either 25 ms (PAS_25_) or 35 ms (PAS_35_), and the order of implementation was semi-randomly determined and counterbalanced across participants. During stimulation, participants were seated reclined with arms relaxed and supported by a pillow, and their eyes were open (Fig. 1a). They wore earplugs to protect their ears from clicking sounds generated by TMS^38^. Before and after the PAS sessions, EEG measurements were also separately taken with ES alone and TMS alone (Fig. 1b).

PAS was examined according to established methods^1,2,10,11^. During each PAS session, 200 stimulus pairs were applied with an average interval of 4.2 s (approximately 0.24 Hz, jittered, 4.0–4.4 s) between stimulus pairs. Each PAS session lasted approximately 15 min. We instructed the participants to pay attention to the ES, relax their whole body, and look at a fixation point (cross marks) attached to the eye position in the natural posture during stimulation.

During ES sessions, 300 stimuli were delivered with an average interval of 525 ms (approximately 1.9 Hz, jittered, 500–550 ms). The relatively high number of 300 stimuli was selected to obtain clean somatosensory-evoked potential (SEP) waveforms with minimal noise, facilitating waveform comparisons before and after PAS. Each ES session lasted approximately 3 min. During TMS sessions, 30 stimuli were delivered with an average interval of 5.25 s (approximately 0.19 Hz, jittered, 5.0–5.5 s). The TMS session was placed between the pre- and post-PAS ES sessions, with a lower number of stimuli (30) chosen to minimize the delay of the post-PAS ES measurements relative to the PAS session. Each TMS session lasted approximately 3 min.

We installed the PAS and EEG settings in an electromagnetically shielded and soundproof room. An EEG recording computer, a stimulation control computer, and a constant-current stimulator (DS8R, Digitimer Ltd., Herts, UK) were placed outside the shielded room. The computer for stimulus control simultaneously controlled the constant-current stimulator and TMS device in the shielded room.

### EEG, electromyography (EMG), and electrooculography (EOG) recordings

A TMS-compatible DC-EEG amplifier (BrainAmp MR plus, BrainProducts GmbH, Germany) was used. EEG was recorded from 63 scalp sites using 2 mm thick, flat Ag/AgCl TMS-compatible C-ring electrodes with a slit to minimize TMS-induced eddy currents in a closed ring, referenced to the right earlobe with ground at AFz, and mounted on a 10/10 system EasyCap (EASYCAP GmbH, Germany)^18^. The electrode sites included Fpz, Fz, FCz, Cz, CPz, Pz, POz, Oz, Iz, Fp1, Fp2, AF3, AF4, AF7, AF8, F1–F8, FC1–FC8, C1–C8, CP1–CP8, P1–P8, PO3, PO4, PO7, PO8, O1, O2, O9, and O10. EEG signals were continuously recorded at a sampling rate of 5 kHz with an online filter between DC and 1,000 Hz and a resolution of 0.1 µV. Whenever possible, impedance was kept below 5 kΩ (and always below 20 kΩ). The electrode lead wires were arranged orthogonal to the TMS coil handle direction to reduce TMS-induced artifacts^38^.

The signals were simultaneously recorded by EMG of the first dorsal interosseous (FDI) muscle with surface electrodes (Covidien Commercial Ltd., UK) using a belly-tendon montage. The ground was positioned at the right olecranon. These signals were recorded at DC with a sampling rate of 1 kHz and a resolution of 0.5 µV. Horizontal and vertical EOG were also measured.

### TMS

Monophasic TMS pulses were delivered to the hand area of the left M1 using a MagVenture MagPro X100 stimulator with a figure-8 coil (MC-B70, MagVenture Inc., Denmark). The recharge delay was set to 2,000 ms to avoid including recharging artifacts in the data around TMS onset. The coil was held tangentially to the skull. The handle pointed backward perpendicular to the assumed line of the central sulcus, inducing posterior-to-anterior electric current in the brain.

The stimulation site and intensity were selected by the following procedure. First, the M1 hot spot that evokes the largest MEP for the right FDI muscle was determined. Second, the individual intensity of a resting motor threshold that induces a MEP of > 50 µV from the right FDI muscle in five out of ten trials was determined. During all sessions, the TMS stimulus intensity was selected so as to evoke a peak-to-peak MEP of 1 mV in the relaxed FDI muscle but to not exceed 120% of the resting motor threshold^2,10^. Coil position and orientation were constantly monitored using the coil navigator Brainsight (Rogue Research Inc., Canada).

### Electrical stimulation

Peripheral stimulation was delivered by a constant-current stimulator (DS8R Digitimer Ltd., United Kingdom) using square-wave pulses with a pulse width of 200 µS. Surface electrodes (GE Healthcare, USA; 8 mm in diameter, 30 mm apart) were fixed with conductive paste to the right wrist over the ulnar nerve. The stimulus intensity was set to a value that does not cause discomfort to participants, with the upper limit being three times that of the sensory threshold.

## Data records

Raw and preprocessed EEG data are available in the accompanying data repository CBS Data Sharing Platform (Shikauchi Y., and Kitajo K., https://doi.org/10.60178/cbs.20240220-001, Creative Commons Attribution-NonCommercial-NoDerivatives 4.0 International (CC BY-NC-ND 4.0)), which includes a detailed data descriptor that provides additional information. It includes raw data from nine participants and pre-processed data from seven participants with sufficient data quality. This dataset was stored in accordance with the brain imaging data structure (BIDS) format^39^. The dataset comprises the following contents.

### Raw data

EEG, EOG, and EMG data are saved in the BrainVision Data Exchange Core Format 1.0 (.eeg, .vhdr, and .vmrk). For each day, there are two files: 1) a series of measurements from the pre-PAS ES, pre-PAS TMS and PAS sessions, named run-01_eeg; 2) a series of measurements from the post-PAS TMS and post-PAS ES sessions, named run-02_eeg. This separation allows the data handler to analyze the data after PAS without knowing the PAS conditions (i.e., triple-blind). The Brainsight data on the TMS coil location were saved as tab-delimited text format (*_brainsight-motion.tsv).

### Preprocessed data

All preprocessed data for each participant were saved in Mat file format (.mat) (see Technical Validation). Demographic information and stimulus parameters for each participant were stored in tab-delimited text format.

## Technical Validation

This section presents the analyses that were performed to ensure the quality of EEG measurements and stimulation techniques. First, we show the reliability of the coil location to demonstrate that TMS was stably applied to the target site. Next, the stimulus-evoked potentials in the ES and TMS sessions show the quality of the EEG measurements. Lastly, MEP and TEP are shown to illustrate the effect of PAS and quality of EEG measurements during PAS.

### TMS coil positioning

We performed TMS while adjusting the coil position according to the magnetic resonance (MR) image-guided navigation system, which measured the error between the estimated stimulation site on the cortex from the real-time measured coil position and target site (e.g., hot spot and target error; Fig. 3). Additionally, the angle of the coil’s stimulating surface along the head (angle error) and the rotation of the coil’s stimulating surface (twist error) were measured to determine how many degrees they deviate from the posture determined when the resting motor threshold was calculated. The median and 25^th^ and 75^th^ percentiles of the errors recorded by the system were obtained offline, session by session (Fig. 3). Trials in which measurements failed due to head movements of the participants were excluded from the analysis. In the EEG and EMG analyses that followed, no other procedures were performed based on this measurement system.

### EEG data processing

We used MATLAB (Mathworks, USA) scripts that were developed in-house with the FieldTrip toolbox^40^ for data processing. All EEG data were re-referenced offline to the average left and right earlobe electrode signals and then processed in different steps, depending on the session types.

### SEPs in the ES sessions

The SEP waveforms were obtained from the re-referenced data during ES sessions using the following procedure: First, EEG data were applied to a 3 Hz high-pass filter. Second, the data were epoched from −150 to 300 ms intervals around the onset of the ES pulse. Third, we discarded epochs within pre- or post-stimulation (−150–0 ms and 50–300 ms) in which the EEG amplitude exceeded ± 150 µV. On average, 269 (range 202–300) trials remained in each session. Finally, trial averages were obtained for each session.

The average waveform confirmed N20, P25, N30, and P45, which have often been reported as SEP with upper limb ES^41,42^. Moreover, the topography of those components was similar across the four conditions (Fig. 4). These results suggest that ES was stable regardless of the experimental conditions.

### TEPs in the TMS sessions

To obtain the grand-average TEP, following the procedure shown in Fig. 2, EEG data were epoched from −300 to 1,000 ms intervals around the onset of the TMS pulse. Referring to previously reported methods^27,43,44^, the following procedure was used to remove TMS artifacts and noisy epochs: First, a period of the TMS onset to 20 ms was linearly interpolated, as this is the time frame during which the ringing artifact can be visually observed. Second, we discarded epochs within pre- or post-stimulation (−300 to 0 ms and 50 to 300 ms) in which the EEG amplitude exceeded ± 150 µV with a baseline of −200 to −5 ms. On average, 30 (range 28–30) trials remained in each session. Third, we attenuated two types of artifacts using independent component analysis^45^. Independent components with extremely large amplitudes, i.e., those with maximum z-score values of > 6 between 0 and 50 ms and with a correlation with EOG of ≥ 0.1 were removed^46^. On average, 12 (range 4–20) components remained in each session. Fourth, a period of the TMS onset to 20 ms was linearly interpolated. Finally, a second-order, Butterworth, zero-phase band-pass filter (1–80 Hz) was used for the trend removals (Fig. 5).

**Figure 2.**
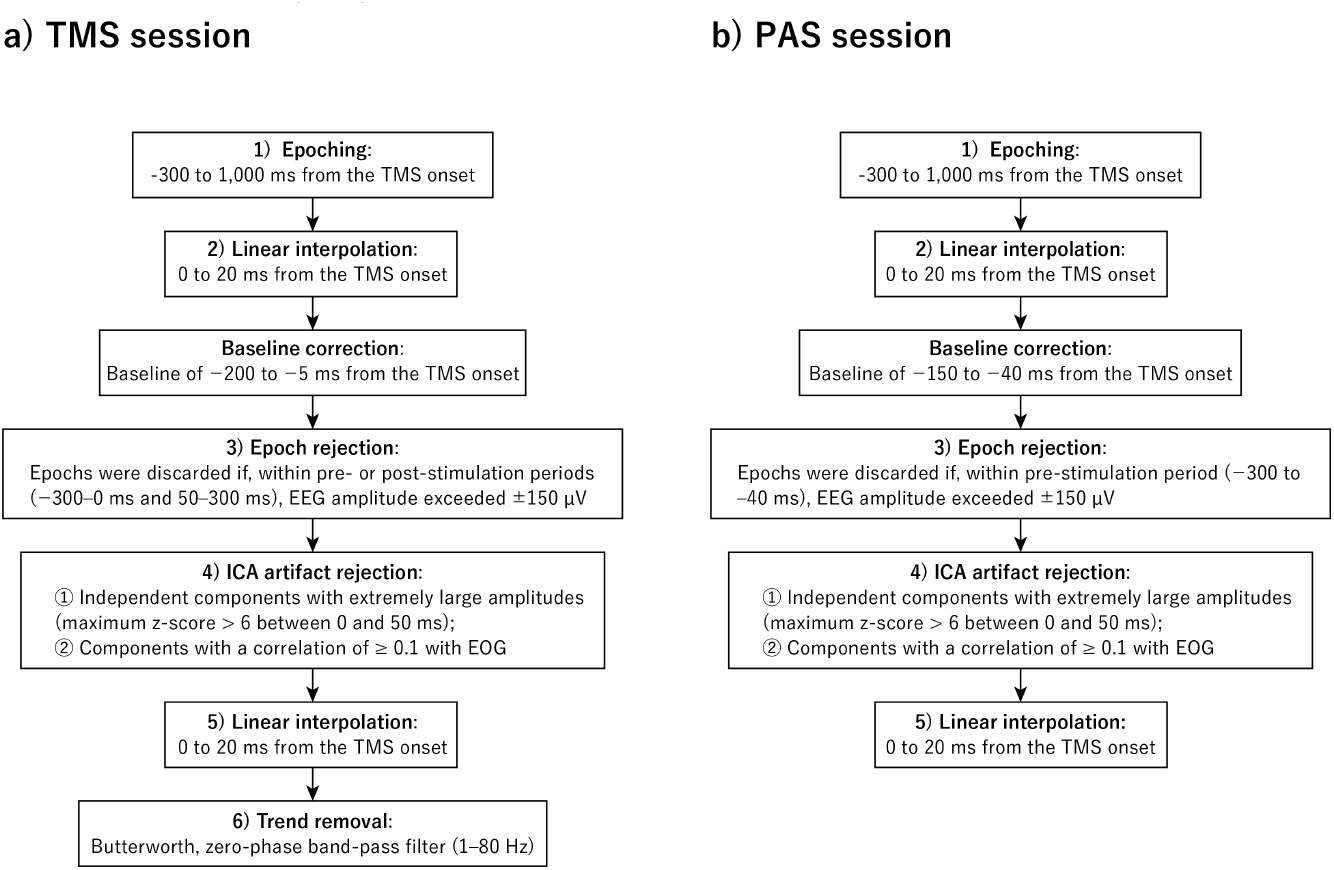
EEG Preprocessing workflow for TMS session (a) and PAS session (b).

**Figure 3.**
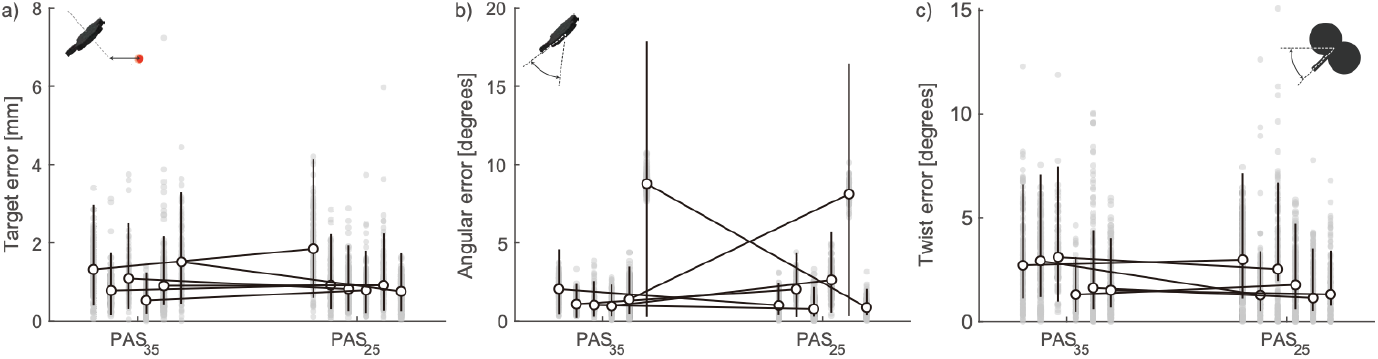
Variations in coil posture during paired associative stimulation (PAS) sessions. The open circles indicate the medians of each participant in the PAS sessions, the upper and lower whiskers indicate the 75th and 25th percentiles, respectively, and the gray dots indicate the values obtained for each trial. a) The difference between the transcranial magnetic stimulation target site on the cortex (i.e., hot spot, a red area) and the stimulus site estimated from the position and orientation of the coil. b) Errors in the angle of the coil relative to the head surface are caused by the coil floating due to the electroencephalography electrodes (which are about 2 mm thick) and by the changing shape of the head surface in contact with the coil due to shifting of the stimulation position observed in (a). c) The coil twist was set to the angle at which motor-evoked potentials are most easily observed for each individual (i.e., roughly 45 degrees from the midline). The twist error indicates rotation of the coil handle from the defined coil twist.

**Figure 4.**
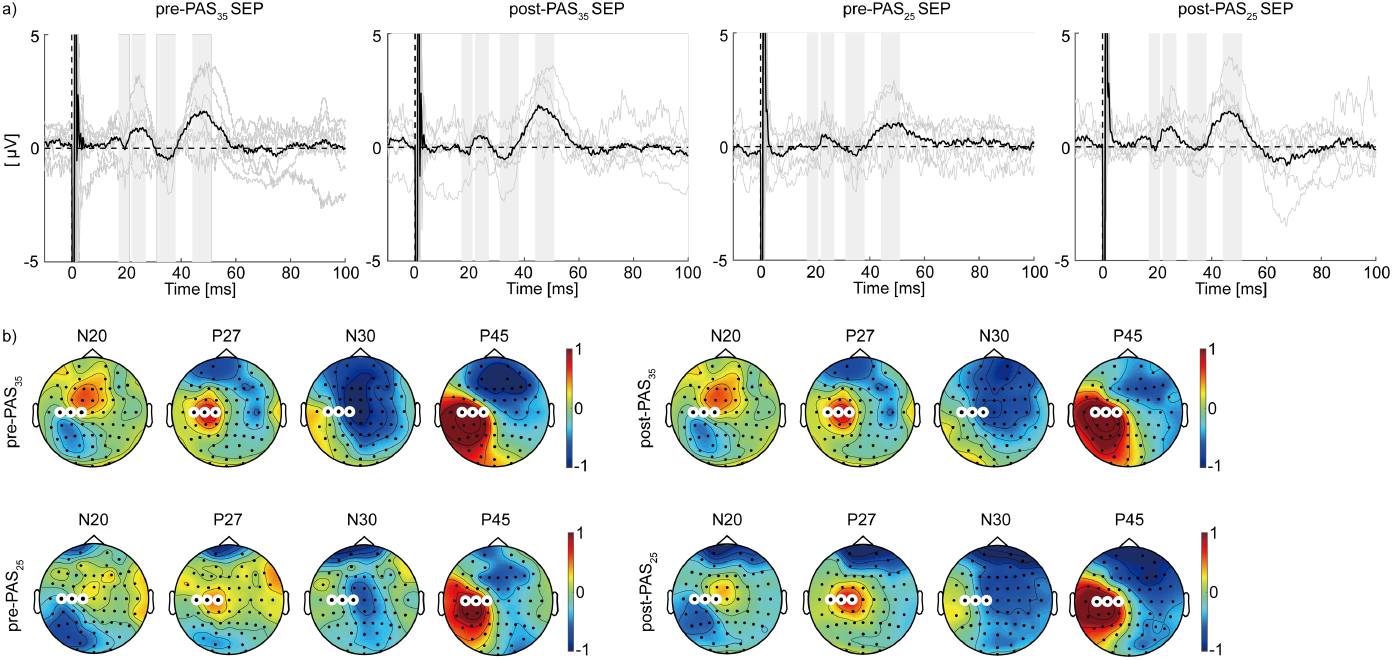
Average waveform and topography of sensory-evoked potential (SEP) to electrical stimuli delivered to the left rest. a) The black line shows the ground average of three channels (C1, C3, and C5) time-locked at the onset of electrical stimulation. The gray lines represent the trial average for individuals. Gray bands indicate the time windows used to determine the topology of N20, P27, N30, and P45 in (b). b) Channels highlighted with white lines are C1, C3, and C5.

**Figure 5.**
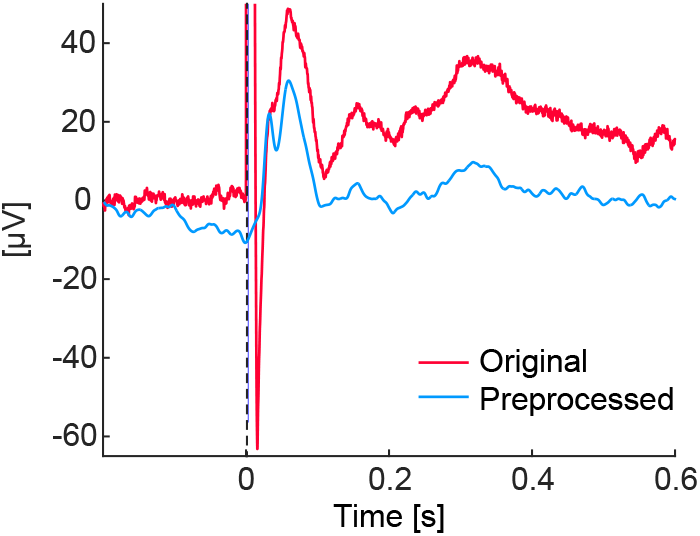
Transcranial magnetic stimulation artifact removal. TMS-evoked potential data before and after artifact removal, consisting of 30 trial averages of C3 signals for a representative individual (subject 3).

The average waveform confirmed P30, P60, N100, and P180, which have often been reported as TEP with left M1 stimulation^47,48^. Moreover, the topography of those components was similar across the four conditions (Fig. 6). These results suggest that we successfully maintained a consistent level of data quality regardless of experimental conditions.

**Figure 6.**
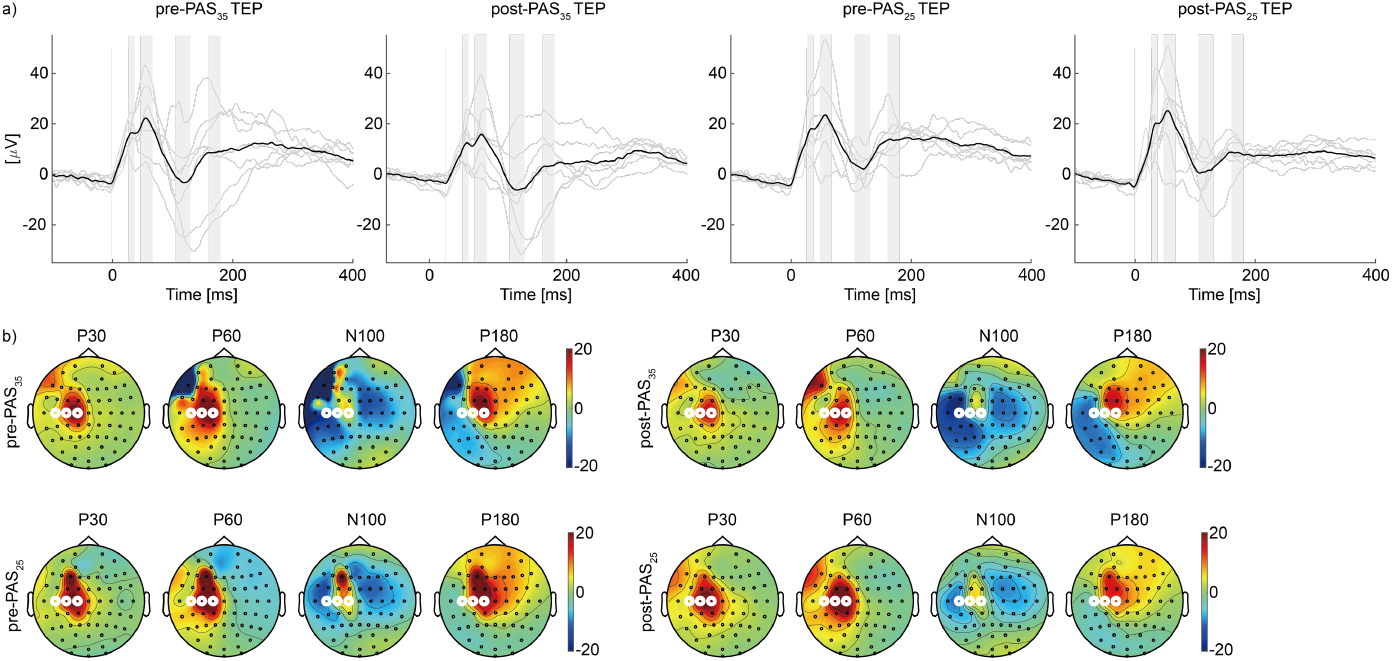
Transcranial magnetic stimulation (TMS)-evoked responses before and after paired associative stimulation. a) The black line shows the ground average of three channels (C1, C3, and C5) time-locked at the onset of TMS. The thin gray lines represent the trial average for an individual. Gray bands indicate the time windows used to determine the topography of P30, P60, N100, and P180 in (b). b) Channels highlighted with white lines are C1, C3, and C5.

### TEPs in the PAS sessions

The TEP waveforms in the PAS sessions were processed according to the same procedure as the TMS sessions (Fig. 2), with the following modifications: we discarded epochs within pre-stimulation (from −300 to −40 ms) in which the EEG amplitude exceeded ± 150 µV with a baseline of −150 to −40 ms (−40 ms was chosen to precede the earliest ES onset at −35 ms). On average, 8 (range 3–13) components remained in each session. Finally, a period of the TMS onset to 20 s was linearly interpolated. TEP during PAS showed a similar temporal and spatial pattern to that during the MEP session (Fig. 8).

### The effects of PAS on MEPs

To measure the effects of PAS_35_ and PAS_25_, the amplitudes of FDI MEPs during pre- and post-PAS MEP sessions were calculated. The peak-to-peak amplitudes of MEPs (mV) were compared within individuals. The trials with background EMG activity more than 1.5 interquartile ranges above the upper quartile or below the lower quartile were rejected to eliminate the effects of increased MEP amplitudes due to voluntary contractions. Post-PAS MEPs were normalized to median pre-PAS MEPs for each condition.

Of the seven participants, three had significantly greater post-PAS_25_ MEP than post-PAS_35_ MEP (p < 0.05, two-sample t-test) (Fig. 7), and six exhibited a trend of higher median MEP amplitudes in the PAS_25_ than in the PAS_35_ condition (one-tailed Wilcoxon signed-rank test, p = 0.055).

**Figure 7.**
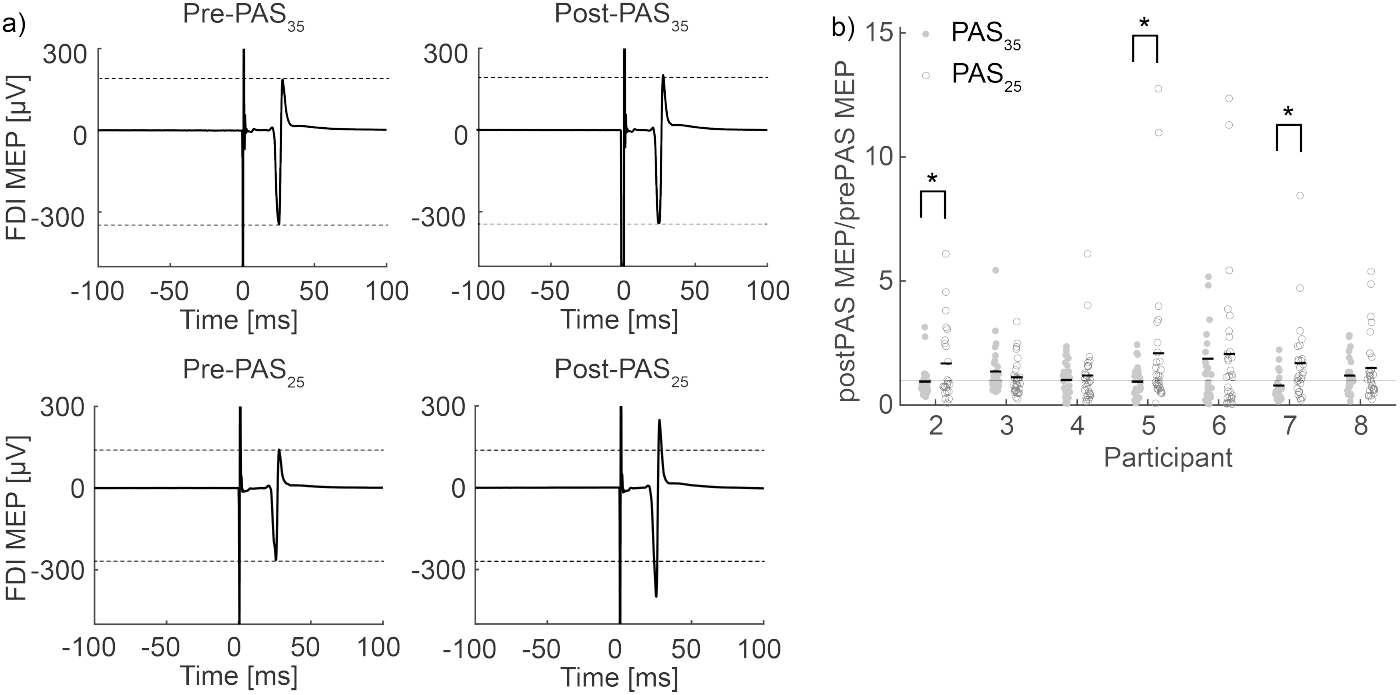
Electromyography (EMG) changes with paired associative stimulation (PAS). a) Trial average of EMG induced by transcranial magnetic stimulation in a representative participant (subject 4). Dotted lines indicate motor-evoked potential (MEP) amplitudes before PAS. b) Comparison of the EMG change under PAS_35_ and PAS_25_. The mean ratio is indicated by a black line. Asterisks (*) indicate significant differences between the two conditions (p < 0.05, two-sample t-test). The gray line delineates the value of 1. FDI: first dorsal interosseous muscle.

**Figure 8.**
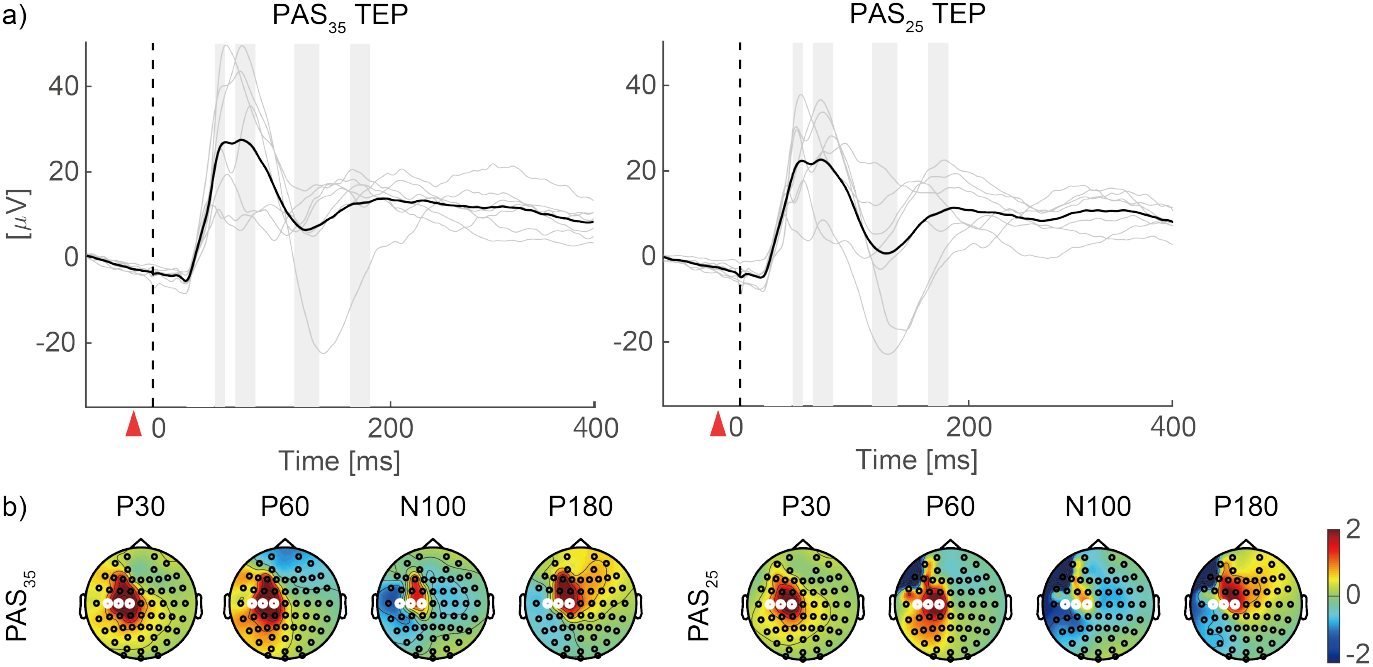
Transcranial magnetic stimulation (TMS)-evoked responses and their topography during paired associative stimulation (PAS). a) The black line shows the ground average of three channels (C1, C3, and C5) time-locked at the onset of TMS. The thin gray lines represent the trial averages for individuals. Red triangles indicate the onset of electrical stimulation. Gray bands indicate the time windows used to determine the topography of P30, P60, N100, and P180 in (b). b) Channels highlighted with white lines are C1, C3, and C5. TEP: TMS-evoked potential.

## Code availability

The code used in this study is available in the data repository CBS Data Sharing Platform (Shikauchi Y., and Kitajo K., https://doi.org/10.60178/cbs.20240220-001, Creative Commons Attribution-NonCommercial-NoDerivatives 4.0 International (CC BY-NC-ND 4.0)) and includes the scripts used for data preprocessing, analysis, and visualization.

## Acknowledgements

We would like to thank Prof. Takashi Hanakawa and Prof. Yoshihiro Noda for their expertise and guidance regarding the safety of our experiment. Prof. Noda and his colleague gave us the opportunity to observe their PAS experiment. We gratefully acknowledge the technical assistance of Ms. Kaori Maeda, Mr. Takuro Naruse, and Ms. Kaho Magami. This project was supported by a search grant from TOYOTA Motor Corporation to K.K. and KAKENHI Grant Numbers JP18K15341, 19J00733, and 21K19810 (all to Y.S.).

## Author contributions

Study design: Y.S. and K.K.; Experimental plan: Y.S. and K.U.; Data acquisition: Y.S.; Data analysis plan: Y.S., K.U., Y.O.O., and K.K.; Data analysis: Y.S.; Drafting the article: Y.S.; Additions and critical revisions: K.U., Y.O.O., and K.K.

## References

1. Suppa, A. et al. The associative brain at work: Evidence from paired associative stimulation studies in humans. Clin. Neurophysiol. 128, 2140–2164 (2017).

2. Stefan, K., Kunesch, E., Cohen, L. G., Benecke, R. & Classen, J. Induction of plasticity in the human motor cortex by paired associative stimulation. Brain 123, 572–584 (2000).

3. Alder, G., Signal, N., Olsen, S. & Taylor, D. A Systematic Review of Paired Associative Stimulation (PAS) to Modulate Lower Limb Corticomotor Excitability : Implications for Stimulation Parameter Selection and Experimental Design. 13, (2019).

4. Sasaki, T. et al. Modulation of motor learning by a paired associative stimulation protocol inducing LTD-like effects. Brain Stimul. 11, 1314–1321 (2018).

5. Hooyman, A., Garbin, A., Fisher, B. E., Kutch, J. J. & Winstein, C. J. Paired associative stimulation applied to the cortex can increase resting-state functional connectivity: A proof of principle study. Neurosci. Lett. 784, 136753 (2022).

6. Fiori, F., Chiappini, E. & Avenanti, A. Enhanced action performance following TMS manipulation of associative plasticity in ventral premotor-motor pathway. Neuroimage 183, 847–858 (2018).

7. Hernandez-Pavon, J. C., San Agustín, A., Wang, M. C., Veniero, D. & Pons, J. L. Can we manipulate brain connectivity? A systematic review of cortico-cortical paired associative stimulation effects. Clin. Neurophysiol. 154, 169–193 (2023).

8. Müller-Dahlhaus, J. F. M., Orekhov, Y., Liu, Y. & Ziemann, U. Interindividual variability and age-dependency of motor cortical plasticity induced by paired associative stimulation. Exp. Brain Res. 187, 467–475 (2008).

9. López-Alonso, V., Cheeran, B., Río-Rodríguez, D. & Fernández-Del-Olmo, M. Inter-individual variability in response to non-invasive brain stimulation paradigms. Brain Stimul. 7, 372–380 (2014).

10. Player, M. J., Taylor, J. L., Alonzo, A. & Loo, C. K. Paired associative stimulation increases motor cortex excitability more effectively than theta-burst stimulation. Clin. Neurophysiol. 123, 2220–2226 (2012).

11. Classen, J. et al. Chapter 59 Paired associative stimulation. Suppl. Clin. Neurophysiol. 57, 563–569 (2004).

12. Siebner, H. R. et al. Transcranial magnetic stimulation of the brain: What is stimulated? - A consensus and critical position paper. Clin. Neurophysiol. 140, 59–97 (2022).

13. Kasai, T., Kawai, S., Kawanishi, M. & Yahagi, S. Evidence for facilitation of motor evoked potentials (MEPs) induced by motor imagery. Brain Res. 744, 147–150 (1997).

14. Bergmann, T. O. et al. EEG-guided transcranial magnetic stimulation reveals rapid shifts in motor cortical excitability during the human sleep slow oscillation. J. Neurosci. 32, 243– 253 (2012).

15. Silvanto, J. & Pascual-Leone, A. State-dependency of transcranial magnetic stimulation. Brain Topogr. 21, 1–10 (2008).

16. Conde, V. et al. Cortical thickness in primary sensorimotor cortex influences the effectiveness of paired associative stimulation. Neuroimage 60, 864–870 (2012).

17. Murase, N., Cengiz, B. & Rothwell, J. C. Inter-individual variation in the after-effect of paired associative stimulation can be predicted from short-interval intracortical inhibition with the threshold tracking method. Brain Stimul. 8, 105–113 (2015).

18. Hernandez-Pavon, J. C. et al. TMS combined with EEG: Recommendations and open issues for data collection and analysis. Brain Stimul. 16, 567–593 (2023).

19. Belardinelli, P. et al. Reproducibility in TMS-EEG studies: A call for data sharing, standard procedures and effective experimental control. Brain stimulation vol. 12 787–790 (2019).

20. Tremblay, S. et al. Clinical utility and prospective of TMS–EEG. Clin. Neurophysiol. 130, 802–844 (2019).

21. Arrigoni, E., Bolognini, N., Pisoni, A. & Guidali, G. Neurophysiological correlates of plasticity induced by paired associative stimulation (PAS) targeting the motor cortex: a TMS-EEG registered report. (2023).

22. Stokes, M. G. et al. Simple metric for scaling motor threshold based on scalp-cortex distance: application to studies using transcranial magnetic stimulation. J. Neurophysiol. 94, 4520–4527 (2005).

23. Sommer, M. et al. Half sine, monophasic and biphasic transcranial magnetic stimulation of the human motor cortex. Clin. Neurophysiol. 117, 838–844 (2006).

24. Hernandez-Pavon, J. C., Kugiumtzis, D., Zrenner, C., Kimiskidis, V. K. & Metsomaa, J. Removing artifacts from TMS-evoked EEG: A methods review and a unifying theoretical framework. J. Neurosci. Methods 376, 109591 (2022).

25. Veniero, D., Bortoletto, M. & Miniussi, C. TMS-EEG co-registration: on TMS-induced artifact. Clin. Neurophysiol. 120, 1392–1399 (2009).

26. Mutanen, T., Mäki, H. & Ilmoniemi, R. J. The effect of stimulus parameters on TMS-EEG muscle artifacts. Brain Stimul. 6, 371–376 (2013).

27. Rogasch, N. C. et al. Removing artefacts from TMS-EEG recordings using independent component analysis: importance for assessing prefrontal and motor cortex network properties. Neuroimage 101, 425–439 (2014).

28. ter Braack, E.M., de Vos, C. C. & van Putten M.J.A.M. Masking the Auditory Evoked Potential in TMS-EEG: A Comparison of Various Methods. Brain Topogr. 28, 520–528 (2015).

29. Russo, S. et al. TAAC - TMS Adaptable Auditory Control: A universal tool to mask TMS clicks. J. Neurosci. Methods 370, 109491 (2022).

30. Biabani, M., Fornito, A., Mutanen, T. P., Morrow, J. & Rogasch, N. C. Characterizing and minimizing the contribution of sensory inputs to TMS-evoked potentials. Brain Stimul. 12, 1537–1552 (2019).

31. Rocchi, L. et al. Disentangling EEG responses to TMS due to cortical and peripheral activations. Brain Stimul. 14, 4–18 (2021).

32. Rossi, S., Hallett, M., Rossini, P. M. & Pascual-Leone, A. Safety, ethical considerations, and application guidelines for the use of transcranial magnetic stimulation in clinical practice and research. Clin. Neurophysiol. 120, 2008–2039 (2009).

33. Rossini, P. M. et al. Non-invasive electrical and magnetic stimulation of the brain, spinal cord, roots and peripheral nerves: Basic principles and procedures for routine clinical and research application. An updated report from an I.F.C.N. Committee. Clin. Neurophysiol. 126, 1071–1107 (2015).

34. Oldfield, R. C. The assessment and analysis of handedness: The Edinburgh inventory. Neuropsychologia 9, 97–113 (1971).

35. Bikson, M. et al. Guidelines for TMS/tES clinical services and research through the COVID-19 pandemic. Brain Stimulation: Basic, Translational, and Clinical Research in Neuromodulation 13, 1124–1149 (2020).

36. Sale, M. V., Ridding, M. C. & Nordstrom, M. A. Factors influencing the magnitude and reproducibility of corticomotor excitability changes induced by paired associative stimulation. Exp. Brain Res. 181, 615–626 (2007).

37. Lang, N. et al. Circadian modulation of GABA-mediated cortical inhibition. Cereb. Cortex 21, 2299–2306 (2011).

38. Sekiguchi, H., Takeuchi, S., Kadota, H., Kohno, Y. & Nakajima, Y. TMS-induced artifacts on EEG can be reduced by rearrangement of the electrode’s lead wire before recording. Clin. Neurophysiol. 122, 984–990 (2011).

39. Gorgolewski, K. J. et al. The brain imaging data structure, a format for organizing and describing outputs of neuroimaging experiments. Sci Data 3, 160044 (2016).

40. Oostenveld, R., Fries, P., Maris, E. & Schoffelen, J.-M. FieldTrip: open source software for advanced analysis of MEG, EEG, and invasive electrophysiological data. Comput. Intell. Neurosci. 2011: 156869, (2011).

41. Desmedt, J. E., Huy, N. T. & Bourguet, M. The cognitive P40, N60 and P100 components of somatosensory evoked potentials and the earliest electrical signs of sensory processing in man. Electroencephalogr. Clin. Neurophysiol. 56, 272–282 (1983).

42. Arslanova, I., Wang, K., Gomi, H. & Haggard, P. Somatosensory evoked potentials that index lateral inhibition are modulated according to the mode of perceptual processing: comparing or combining multi-digit tactile motion. Cogn. Neurosci. 13, 47–59 (2022).

43. Herring, J. D., Thut, G., Jensen, O. & Bergmann, T. O. Attention modulates TMS-locked alpha oscillations in the visual cortex. J. Neurosci. 35, 14435–14447 (2015).

44. Okazaki, Y. O., Mizuno, Y. & Kitajo, K. Probing dynamical cortical gating of attention with concurrent TMS-EEG. Sci. Rep. 10, 1–10 (2020).

45. Korhonen, R. J. et al. Removal of large muscle artifacts from transcranial magnetic stimulation-evoked EEG by independent component analysis. Med. Biol. Eng. Comput. 49, 397–407 (2011).

46. Okazaki, Y. O., Nakagawa, Y., Mizuno, Y., Hanakawa, T. & Kitajo, K. Frequency- and Area-Specific Phase Entrainment of Intrinsic Cortical Oscillations by Repetitive Transcranial Magnetic Stimulation. Front. Hum. Neurosci. 15, 608947 (2021).

47. Komssi, S., Kähkönen, S. & Ilmoniemi, R. J. The effect of stimulus intensity on brain responses evoked by transcranial magnetic stimulation. Hum. Brain Mapp. 21, 154–164 (2004).

48. Conde, V. et al. The non-transcranial TMS-evoked potential is an inherent source of ambiguity in TMS-EEG studies. Neuroimage 185, 300–312 (2019).

